# Neural correlates of exceptional memory ability in SuperAgers: A multimodal approach using FDG-PET, PIB-PET, and MRI

**DOI:** 10.1101/666438

**Authors:** Wyllians Vendramini Borelli, Eduardo Leal-Conceição, Michele Alberton Andrade, Nathalia Bianchini Esper, Paula Kopschina Feltes, Ricardo Bernardi Soder, Cristina Sebastião Matushita, Louise Mross Hartmann, Graciane Radaelli, Lucas Porcello Schilling, Cristina Moriguchi-Jeckel, Ana Maria Marques da Silva, Mirna Wetters Portuguez, Alexandre Rosa Franco, Jaderson Costa da Costa

## Abstract

Individuals at 80 years of age or above with exceptional memory are considered SuperAgers (SA). A multimodal brain analysis of SA may provide biomarkers of successful cognitive aging. Herein, a molecular (PET-FDG, PET-PIB), functional (fMRI) and structural analysis (MRI) of SA was conducted. Ten SA, ten age-matched older adults (C80) and ten cognitively normal middle-aged adults underwent cognitive testing and neuroimaging examinations. The relationship between cognitive scores and cingulate areas and hippocampus were examined. The SA group showed increased FDG SUVr in the left subgenual Anterior Cingulate Cortex (sACC, p<0.005) as compared to that in the C80 group. Amyloid deposition was similar between SA and C80 in the described regions or overall areas (p>0.05). The SA group also presented decreased connectivity between left sACC and posterior cingulate (p<0.005) as compared to that of C80 group. These results support the key role of ACC in SA, even in the presence of amyloid deposition. It also suggests that sACC can be used as a potential memory biomarker in older adults.

**Abbreviations:** BCa – Bias corrected accelerated: SA – SuperAgers: C50 – Middle-aged controls: C80 – Age-matched controls

## 1. Introduction

SuperAgers (SA) is a definition to describe individuals who have achieved successful cognitive aging (Borelli et al., 2018a). By definition, SA are individuals aged 80 years or more, exhibiting higher youthful memory scores, which are similar to those of younger individuals who are 50 to 65 years old (Harrison et al., 2012). A review that presents the unique brain morphology and function of SA has been recently published (Borelli et al., 2018b). SA show an increased cortical thickness in the anterior cingulate, similar global amyloid burden and lower volumes of white matter hypointensity, as compared to that of normal aged individuals (Baran and Lin, 2018; Harrison et al., 2018). Whether SA activates compensatory mechanisms or they exhibit youthful brain features is a rather controversial topic.

Glucose brain metabolism can be measured by Positron Emission Tomography (PET) using the radiotracer ^18^F-Fluorodeoxyglucose (FDG) (Knopman et al., 2014). This ligand has clinically been used as an important biomarker in Alzheimer’s Disease (AD) (Mosconi et al., 2008; Noble and Scarmeas, 2009) and can be used to measure the decrease in brain activity resulting from normal aging process. Although a reduction in whole-brain glucose metabolism is observed in older adults, the magnitude of decrease differs depending on the region (Nugent et al., 2014). Healthy older adults exhibit significant hypometabolism in the medial frontal regions such as the anterior region of the cingulate, as compared to that in the younger controls (Ishibashi et al., 2018; Pardo et al., 2007; Yoshizawa et al., 2014). A study reported that high-performing older adults exhibited increased prefrontal FDG uptake as compared to their age-matched peers (Cabeza et al., 2002), suggesting that these individuals are less affected by age-related hypometabolism. However, the findings of these participants were not compared to a younger group that would have confirmed whether or not SA exhibited preserved brain metabolism.

Abnormal deposition of amyloid plaques is strongly associated with Alzheimer’s disease spectrum (Jack et al., 2018). A few recent PET studies investigated the amyloid burden in older adults with normal and high cognitive performance using the radiotracer ^11^C-Pittsburgh Compound B (PIB) (Donohue et al., 2017; Jansen et al., 2015; Villemagne et al., 2011). These studies have shown that amyloid-positive older adults exhibited increased rates of cortical atrophy and cognitive decline but decreased regional glucose metabolism in comparison to the amyloid-negative older adults. Older adults with exceptional memory capacity presented mixed findings of brain amyloid load as compared to that of the other normal individuals for that age (Baran and Lin, 2018). However, another follow-up study noted that individuals who maintained their cognitive scores had lower amyloid deposition after three years as compared to individuals whose cognition was declined (Dekhtyar et al., 2017).

Resting-state functional magnetic resonance imaging (rsfMRI) has been shown to be highly accurate in predicting functional connectivity (FC) abnormalities related to the aging process (Iraji et al., 2016). A significant decrease in FC is expected in healthy individuals as they age (Marques et al., 2015), though some brain networks of older adults, involving frontal and temporal areas, are particularly affected (Dennis and Thompson, 2014; Lopez-Larson et al., 2011). The Default Mode Network (DMN), typically associated with memory system and self-thoughts, has been demonstrated to be affected by age-related cognitive decline, thus presenting reduced FC between the anterior cingulate and posterior cingulate in older individuals (Dennis and Thompson, 2014; Koch et al., 2010). Previous studies have highlighted the pivotal role of cingulate regions in successful memory encoding (Grady et al., 2003; Lee et al., 2016) and retrieval (Lega et al., 2017).

Nonetheless, it is not possible to completely evaluate the complexity of human cognitive processes using a single method of analysis. Besides, combining multiple features provided by modality-specific techniques can help improve the overall understanding of the neural basis of successful cognitive aging and SuperAging. Considering the fact that decreased functional activity and altered molecular properties of the brain are expected during the aging process, how these changes relate to each other is still unclear. The aim of the present study was to analyze the changes in the brain in regional glucose metabolism, functional connectivity, amyloid burden and structural characteristics of SA. A multimodal neuroimaging approach (FDG-PET, PIB-PET, functional MRI, and structural MRI) was thus applied to compare SA with a younger, middle-aged control group.

## 2. Methods

### 2.1. Participants

Community-dwelling older and middle-aged adults were evaluated at the Brain Institute of Rio Grande do Sul (BraIns). Individuals were invited via the social media, television advertisement, and also through courses in the university that are focused toward the elderly. The participants included in the study were right-handed individuals with no history of substance abuse, moderate or severe head trauma or serious neurological or psychiatric diseases. All the participants included in the study denied any family history of dementia or cognitive impairment. They also demonstrated preserved daily life activities and negative scores for the Geriatric Depression Scale (raw score < 5) (Yesavage and Sheikh, 1986). Prior to their enrolment into the study, all the participants signed an informed consent form, previously approved by the university’s ethical committee. These participants were then divided in three groups: SuperAgers (SA), Age-matched Controls (C80) and Middle-aged Controls (C50).

The SA group was defined on the basis of previously described criteria (Harrison et al., 2012). This group comprised older adults aged 80 years or above, who showed the ability to a) perform at or above the normative values determined for individuals between 50 to 65 years of age on the delayed-recall score of the Rey Auditory-Verbal Learning Test (RAVLT), and b) perform at or above normative values determined for their age and education in non-memory domains. Non-memory measures included the Mini-Mental State Examination (MMSE), the Trail-Making Test – Part B (TMT-B), the Category Fluency Test–Animals (CFT) and the Boston Naming Test (BNT). Two healthy control groups, comprising age-matched individuals who were running in their 80s (C80) and cognition-matched individuals running in their 50s (C50) were also included in the study. The participants included in both the control groups were required to perform within a normal range in both, memory and non-memory fields, with 1.5 as standard deviation from the mean, using normative values for age and education. A fourth group was selected for comparison purposes, which comprised individuals at a mild stage of Alzheimer’s Disease (AD) following previously established diagnostic criteria (NIA-AA) (McKhann et al., 2011). This group was included to demonstrate extreme, opposite biomarker values compared to the SA and control groups; however, this was not included in the discussion. All the study individuals underwent three imaging sessions, FDG-PET, PIB-PET, and MRI (Fig. 1).

**Figure 1.**
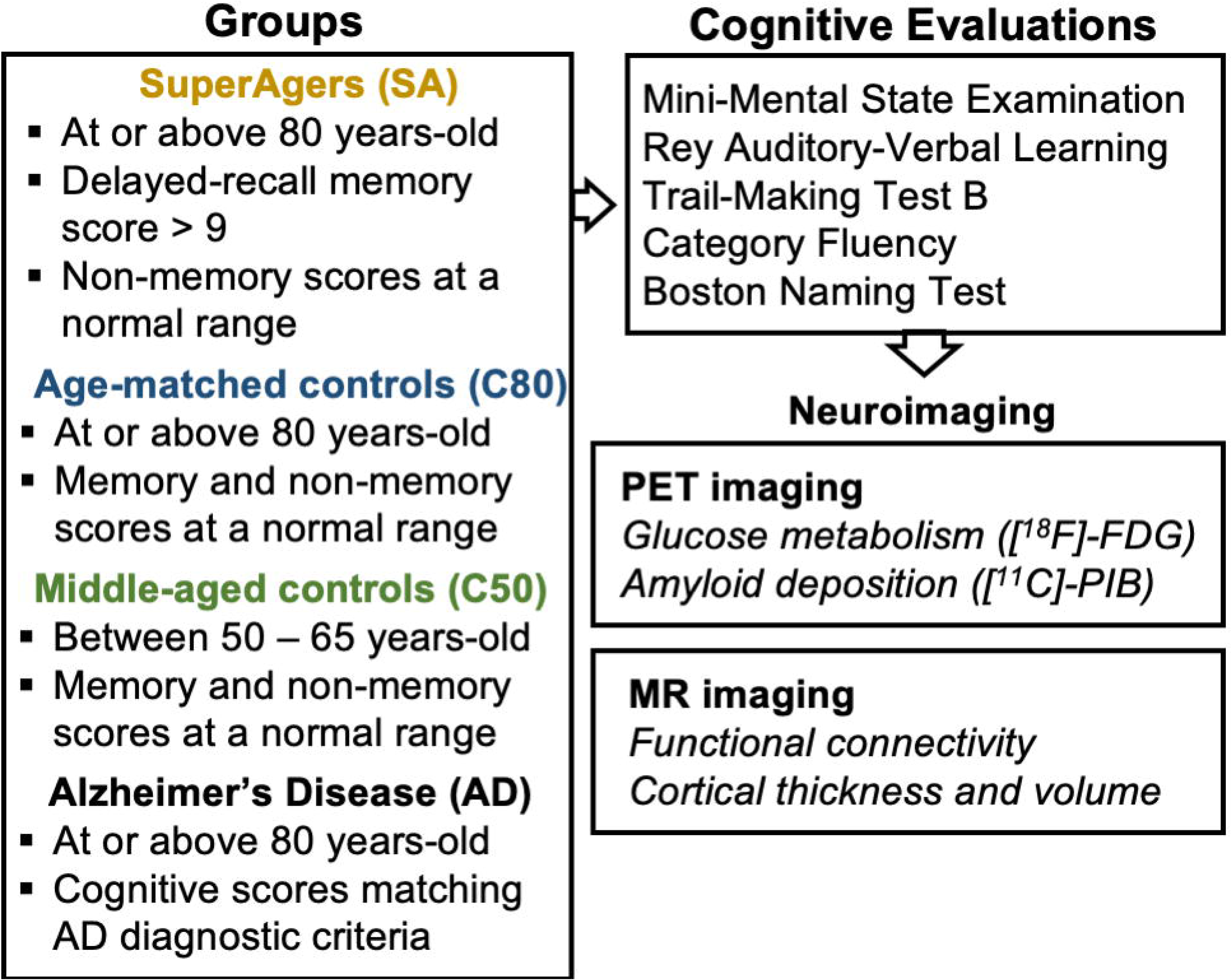
Schematic diagram of the study protocol.

### 2.2. Image acquisition and pre-processing

#### PET imaging

PET scans were performed within five months of clinical screening and cognitive testing. Both FDG and PIB measurements were performed using a GE Discovery 600 scanner. First, a regular CT scan was obtained for attenuation and scatter correction. Subsequently, PET data were acquired using 3D list-mode. The data were reconstructed using VUE Point HD (2 iterations, 32 subsets, filter cutoff 4.8 mm, matrix size 192 × 192 × 47, voxel size 1.56 × 1.56 × 3.27 mm) and corrected for attenuation, scatter, dead time and decay. Before performing PET scans, an intravenous catheter was placed in the left arm of all the patients and their heads were immobilized in order to minimize motion during the scan.

Participants were instructed to follow a low glucose diet 24 hours prior to the FDG scan, in which the last 4-hours consisted of fasting. Before injection, the participants were made to rest for 30 min and capillary glucose was measured 10 min prior to the PET scan, with acceptable levels ranging between 70 and 120 mg/dL, before imaging acquisition. The radiotracer was injected in bolus (326 ± 47 MBq) and a list-mode dynamic emission scan was performed in 60 min. For the PIB scan also, the participants were made to follow a 4-hour fast. PIB was injected intravenously (459 ± 70 MBq) in bolus, while the participants were positioned inside the scanner and a dynamic list-mode acquisition was performed in 90 min.

Static PET images were acquired using raw dynamic frames (6 × 5 min frames) at 30–60 min post-injection for FDG and 40–60 min post-injection for PIB, which were averaged to create one single averaged image. Rigid co-registration of individual data (PET and MRI) was performed using a normalized mutual information algorithm, and the maximum probability atlas (Hammers N30R83) was used for the generation of standard brain regions of interest (ROI) in PNEURO tool (version 3.8, PMOD Technologies Ltd., Zürich, Switzerland).

#### MR imaging

MR structural and functional images were collected on a GE HDxt 3.0T MRI scanner with an 8-channel head coil. A T1-weighted MPRAGE-similar volumetric sequence designed for GE was acquired using 3D FSPGR BRAVO (TR = 6.27ms; TE = 2.25 ms; TI = 550 ms; Flip Angle = 11° matrix size 512 × 512 × 196, voxel size 0.5 × 0.5 × 1.0 mm). These images were used for co-registration with other imaging modalities, tissue segmentation, and definition of ROIs. The echo-planar sequence (EPI) was acquired during a 7-min resting-state protocol (TR/TE=2000/30ms, matrix size 64 × 64, FOV 64 × 64, total volumes 210). During the functional scan, the participants were requested to stare at a crosshair and not to think of anything in particular. A real-time operating system was used to monitor the head motion of the study participants. The participants who moved their heads excessively were reminded to maintain their heads still, and the sequence was restarted.

All functional images were pre-processed using the software AFNI (afni.nimh.nih.gov) (Cox, 1996). Preprocessing steps included slice-time and motion correction and a non-linear spatial normalization to 3.5 × 3.5 × 3.5 mm^3^ voxel template (MNI152 template). Time Repetitions (TR) tracked with excessive motion [Framewise Displacement (FD) > 0.6 mm], were censored from the dataset. The exclusion criterion for the excessive motion was defined to be the motion wherein a participant had 20% of the TRs above the FD threshold. A nuisance regression with six motion estimated parameters (x, y, z, roll, pitch, yaw) and time-series of the average signal of the white matter and cerebrospinal fluid was performed. Signal detrending using a bandpass temporal filter (0.01 and 0.1 Hz) (Weissenbacher et al., 2009) and smoothing with a 6 mm FWHM Gaussian kernel were also employed as preprocessing steps on the functional data.

### 2.3. Neuroimaging Data Analysis

#### PET Imaging

For both FDG and PIB, anatomical ROIs were generated using the Hammers N30R83 brain atlas (Hammers et al., 2003). The Standardized Uptake Value (SUV) was obtained by normalizing tissue concentration to the injected dose and body weight. ROIs were analyzed bilaterally (Gousias et al., 2008; Hammers et al., 2003). The Standardized Uptake Value Ratio (SUVr) was calculated using the cerebellum grey matter as reference (M Bauer et al., 2013), for both FDG and PIB. For PIB images, an adaptation of the AD-signature ROI composite (Jack et al., 2017) was accomplished using the average of the mean uptake in the prefrontal, orbitofrontal, parietal, temporal, anterior and posterior cingulate, and precuneus ROIs. Whole-brain cortical SUVr was measured for FDG analysis. Cingulate regions and hippocampus SUVr were analyzed for both the PET modalities.

#### MR Imaging

##### Seed-based analysis

A seed fMRI (functional MRI) analysis of connectivity was performed, based on previously described regions that distinguished SA from other normal individuals of that age (Baran and Lin, 2018; Harrison et al., 2012). ROIs were extracted by using regional boundaries of the Hammers brain atlas for the following regions: anterior cingulate, posterior cingulate, presubgenual and subgenual areas (Fig. 1: Supplementary material). For each participant, the average time course of the voxels within the seed was collected, and Pearson’s correlation was implemented between the time series of each ROI and all other voxels in the brain. Correlation results were then remodeled using Fisher’s r-to-z method, prior to the statistical analysis.

##### Independent Component Analysis

The data-driven model analysis was also carried out for the rsfMRI with Independent Component Analysis (ICA) using MELODIC (FSL, v6.0.0). Fourteen components were estimated for this analysis, which successfully distinguished resting-state networks; dual regression was applied to identify the independent component (IC) maps in each individual. IC-6 was selected because of significant activation of hippocampal and cingulate areas (Fig. 4).

##### Structural MRI analysis

Cortical thickness and volume were calculated using the image analysis software FreeSurfer (version 6) (Fischl and Dale, 2000). All images were processed by running the “recon-all” script with default settings. Manual corrections were performed for segmentation and parcellation errors, described as a priori by https://surfer.nmr.mgh.harvard.edu/fswiki. This method has been demonstrated to be reliable and was validated with similar accuracy as that of manual segmentation of grey and white matter (Rosas et al., 2002).

### 2.4. Statistical analysis

The normality of the sample was calculated using the Shapiro-Wilk test. ANCOVA and ANOVA calculations followed by Tukey’s post-hoc test were performed to compare SA with other groups for the statistical analysis of FDG-PET and PIB-PET (R Studio – v1.0.136). The seed-based and IC analysis were performed using AFNI’s scripts for group comparison, general linear model fitting (*3dMVM*), and regression analysis (*3dRegAna*) (Cox, 1996). The results were considered to be statistically significant (α<0.05) for a minimum cluster size of 1498 μL and a threshold of p<0.005. The structural MRI analysis (cortical thickness and average volume) was performed using the QDEC interface from Freesurfer. Statistical tests for structural measures were corrected for multiple comparisons using Monte Carlo Simulations (Freesurfer).

Pearson’s correlation tests were employed to measure the relationship between cognitive scores and neuroimaging metrics. Each of these tests used bias-corrected accelerated (BCa) 95% confidence intervals (CI) and 1000 Bootstrapped samples to create a re-sampled range of correlated coefficients. The effect size of all associations was also calculated using Cohen’s d. Multiple linear regressions were performed to calculate the associations between cognitive scores and neuroimaging metrics. All ROI-wise and regression analysis were performed using the R Studio (v1.0.136).

## 3. Results

### 3.1. Demographic factors

No significant differences were found between SA and C80 group in terms of age, years of education or distribution of sex (Table 1). Although C50 group differed from SA group in terms of age, as expected by the inclusion criteria (p < 0.001); yet, education, sex, and cognitive measures did not differ statistically between these groups (p > 0.05 for all measures).

**Table 1.** Demographic Characteristics of the participants

|  | SA | C80 | C50 | AD | SA vs. C80<br>p-values | SA vs. C50<br>p-values |
| --- | --- | --- | --- | --- | --- | --- |
| **Demographic** |  |  |  |  |  |  |
| Age (years) | 82.1 ($\pm 2.5$) | 84.2 ($\pm 3.6$) | 58.5 ($\pm 5.8$) | 84.4 ($\pm 3.6$) | 0.66 | <0.001 |
| Sex (M) | 7 (70%) | 6 (60%) | 8 (80%) | 5 (55.6%) | 0.62 | 0.6 |
| Education<br>(years) | 12.7 ($\pm 4.8$) | 12.9 ($\pm 5.0$) | 14.2 ($\pm 4.9$) | 11.9 ($\pm 5.3$) | 0.99 | 0.91 |
| **Cognitive scores** |  |  |  |  |  |  |
| MMSE | 28.7 ($\pm 1.3$) | 27.9 ($\pm 1.2$) | 29.5 ($\pm 0.7$) | 19.4 ($\pm 5.4$) | 0.91 | 0.91 |
| Delayed-recall scores | 11.4 ($\pm 2.0$) | 6.9 ($\pm 1.6$) | 10.7 ($\pm 3.4$) | 0.3 ($\pm 0.5$) | **0.0002** | 0.86 |
| BNT | 57.5 ($\pm 1.8$) | 54.5 ($\pm 2.7$) | 56.4 ($\pm 3.4$) | 47.9 ($\pm 8.8$) | 0.51 | 0.95 |
| CFT | 20.5 ($\pm 3.6$) | 16.8 ($\pm 8.9$) | 19.5 ($\pm 3.1$) | 10.4 ($\pm 3.0$) | 0.41 | 0.97 |
| TMT-B | 123.3<br>($\pm 40.2$) | 146.9<br>($\pm 64.4$) | 93.3 ($\pm 55.2$) | 278.1<br>($\pm 65.7$) | 0.79 | 0.64 |

SA: SuperAgers; C80: Control–80 years; C50: Controls–50 to 65 years; AD: Alzheimer’s Disease;
MMSE: Mini-mental state examination; BNT: Boston Naming Test; CFT: Category Fluency Test;
TMT-B: Trail-Making Test – Part B.

### 3.2. PET results

#### Brain glucose metabolism

SA group showed significantly increased metabolic activity in sub-regions of the Anterior Cingulate Cortex (ACC), especially the left subgenual ACC (sACC, Table 2), as compared to that in C80 group. The right pre-subgenual region of the ACC (pACC) showed increased brain activity in SA as compared to that in C80 group; however, this relation was later corrected for multiple comparisons (1.02 ±0.08 vs. 0.88 ±0.07, p < 0.01 uncorrected) and did not survive long. Subsequently, sACC and pACC were grouped in a single ROI named ventral anterior cingulate (vACC, Fig. 1 Sup. Material). The SA group showed increased FDG SUVr for both, right and left hippocampal regions as compared to the C80 group (Table 2) but not with C50 group (p > 0.05). These tests did not pass corrections for multiple comparisons.

**Table 2.** Whole-brain and regional metabolic activities across the groups.

|  | SA | C80 | C50 | AD | SA vs. C80<br>p-values |
| --- | --- | --- | --- | --- | --- |
| **FDG-SUV<sub>r</sub>** |  |  |  |  |  |
| Whole-brain cortical uptake | 1.03 ($\pm 0.06$) | 0.97 ($\pm 0.04$) | 1.07 ($\pm 0.06$) | 0.9 ($\pm 0.03$) | 0.18 |
| Right ACC | 0.89 ($\pm 0.14$) | 0.89 ($\pm 0.08$) | 1.04 ($\pm 0.10$) | 0.85 ($\pm 0.08$) | 0.99 |
| Left ACC | 0.87 ($\pm 0.14$) | 0.88 ($\pm 0.12$) | 1.00 ($\pm 0.09$) | 0.87 ($\pm 0.07$) | 0.98 |
| Right PCC | 1.19 ($\pm 0.21$) | 1.16 ($\pm 0.14$) | 1.29 ($\pm 0.11$) | 0.96 ($\pm 0.10$) | 0.95 |
| Left PCC | 1.18 ($\pm 0.21$) | 1.15 ($\pm 0.10$) | 1.28 ($\pm 0.12$) | 0.95 ($\pm 0.09$) | 0.97 |
| Right pACC | 1.02 ($\pm 0.08$) | 0.88 ($\pm 0.07$) | 1.12 ($\pm 0.14$) | 0.86 ($\pm 0.08$) | 0.01* |
| Left pACC | 1.04 ($\pm 0.09$) | 0.93 ($\pm 0.08$) | 1.17 ($\pm 0.10$) | 0.89 ($\pm 0.09$) | 0.04* |
| Right sACC | 0.97 ($\pm 0.08$) | 0.83 ($\pm 0.06$) | 1.05 ($\pm 0.12$) | 0.80 ($\pm 0.10$) | 0.009* |
| Left sACC | 0.99 ($\pm 0.08$) | 0.85 ($\pm 0.05$) | 1.06 ($\pm 0.09$) | 0.81 ($\pm 0.09$) | **0.003\*\*** |
| Right vACC | 0.99 ($\pm 0.07$) | 0.85 ($\pm 0.06$) | 1.08 ($\pm 0.13$) | 1.72 ($\pm 0.56$) | 0.006* |
| Left vACC | 1.02 ($\pm 0.08$) | 0.89 ($\pm 0.06$) | 1.12 ($\pm 0.09$) | 1.80 ($\pm 0.60$) | 0.007* |
| Right Hippocampus | 0.92 ($\pm 0.04$) | 0.81 ($\pm 0.06$) | 0.96 ($\pm 0.05$) | 0.71 ($\pm 0.1$) | 0.006* |
| Left Hippocampus | 0.91 ($\pm 0.05$) | 0.81 ($\pm 0.06$) | 0.97 ($\pm 0.06$) | 0.65 ($\pm 0.09$) | 0.01* |

\*p-values<0.05 uncorrected; \*\*p-values <0.05 after corrected for multiple comparisons. ACC – Anterior Cingulate Cortex. sACC – Subgenual ACC. pACC – Presubgenual ACC. vACC – Ventral ACC.

Importantly, all described regions of the anterior cingulate cortex (including the vACC) and hippocampus exhibited similar metabolic activity between the SA and C50 groups (p > 0.05). In comparison to SA, the AD group showed statistically decreased FDG SUVr for all measures, except for the left ACC, right ACC, left vACC, and right vACC (p > 0.05; Table 2). Besides, ROIs were extracted and compared between the groups. Total volumes of all described regions were observed to be statistically similar between SA and C80 groups (p > 0.05, Table 1 Supplementary data).

Left sACC FDG SUVr further showed a moderate correlation with delayed-recall memory scores for the whole sample (r = 0.41, p < 0.05; BCa 95% CI: r = 0.12:0.62) and for older adults when grouped (SA and C80 groups, r = 0.48, p < 0.05; BCa 95% CI: r = 0.05:0.67) (Fig. 2a). The effect size of this relation was also found to be strong (Cohen’s d = 3.97). Right pACC FDG SUVr revealed a moderate interaction with delayed-recall memory scores for the whole sample (r = 0.59, p = 0.0006; BCa 95% CI: r = 0.3:0.76), and even stronger interaction for the group of older adults (SA and C80 groups, r = 0.71, p < 0.0001; BCa 95% CI: r = 0.38:0.9). A regression model was employed and revealed a strong relationship between right pACC FDG SUVr and delayed-recall scores (p = 0.03) in a model that also accounted for volume and age (overall model R-squared = 0.63, p < 0.001). Though right vACC FDG SUVr exhibited a moderate correlation with delayed-recall memory scores for the entire sample (r = 0.55, p = 0.001; BCa 95% CI: r = 0.22:0.73); however, it depicted a stronger interaction for the group of older adults (SA and C80 groups, r = 0.68, p = 0.0002; BCa 95% CI: r = 0.45:0.78). Right vACC FDG SUVr also showed a significantly positive correlation with the sum of 5 scores of the RAVLT (r = 0.65, p < 0.0001; BCa 95% CI: r = 0.46:0.79), which is usually associated with learning process. After adjustments were made for age and volume, the model showed a significant association between right vACC FDG SUVr and both, delayed-recall scores (overall model R-squared = 0.60, p < 0.0001) and learning scores (overall model R-squared = 0.66, p < 0.0001).

**Figure 2.**
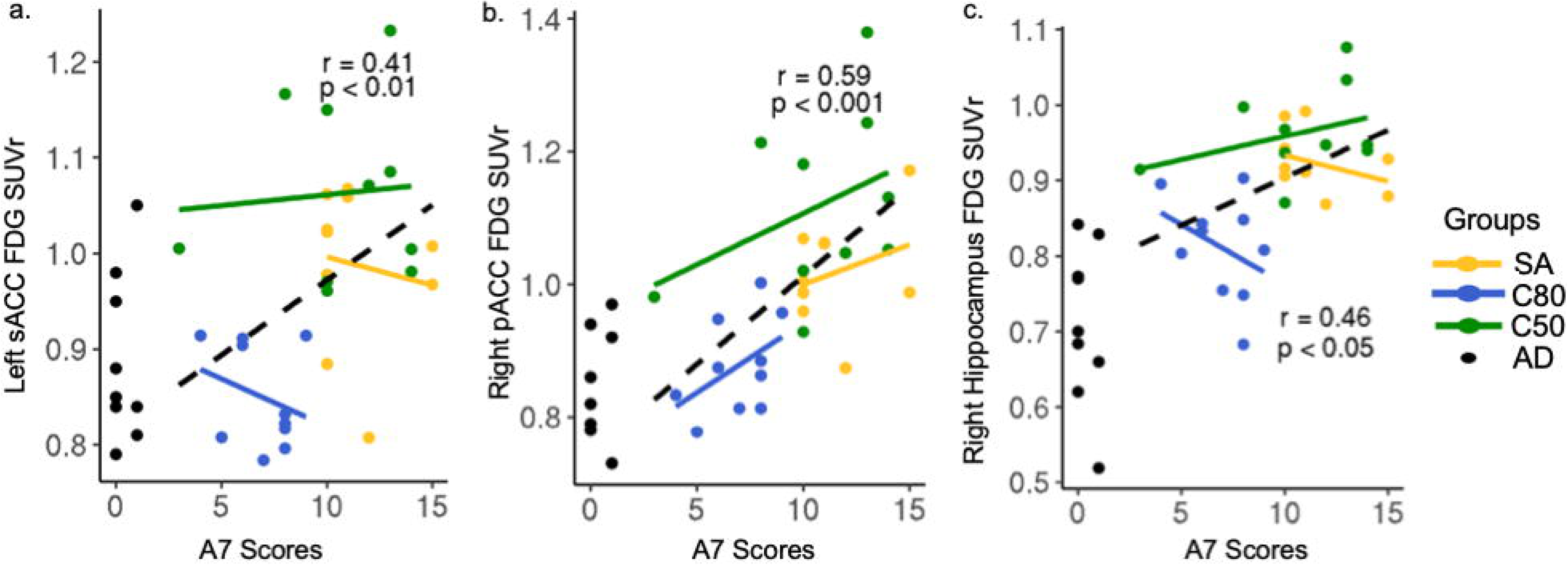
Correlation analysis between ROIs of FDG-PET and, cognitive scores and age. The relation between (a) left sACC FDG SUVr and delayed-recall memory scores, (b) right pACC FDG SUVr and delayed-recall memory scores, (c) right hippocampal FDG SUVr and delayed-recall memory scores. The dashed line represents the whole sample, except the AD group (black dots); while the solid lines represent group tendencies. sACC – subgenual anterior cingulate cortex. pACC – presubgenual anterior cingulate cortex. SUVr – Standardized Uptake Value ratio. SA – SuperAgers. C50 – Middle-aged controls. C80 – Age-matched controls. AD – Alzheimer’s disease.

A moderate correlation was found between both, right and left hippocampal FDG SUVr and the delayed-recall memory scores in the entire sample (r = 0.46 and r = 0.47, p = 0.01 and p = 0.009, respectively) (Fig. 2c). When calculated for each group, the correlations did not reach statistical significance for SA (r = –0.34), C50 (r = 0.35) or C80 (r = –0.36) groups (p > 0.05 for all measures). Though the model exhibited statistical significance (overall model R-squared = 0.39, p < 0.005), there was no association between right (p = 0.09) or left (p = 0.09) hippocampal FDG SUVr in regression models with delayed-recall memory scores when accounted for region volume and age. However, an association between right hippocampal FDG SUVr (p = 0.04) and learning scores was established (overall R-squared = 0.49, p = 0.0004).

A negative correlation between right hippocampal FDG SUVr and age in the whole sample was detected (r = –0.57, p = 0.001; BCa 95% CI: r = –0.75: –0.31), whereas, the SA group alone showed a significantly positive relationship with age (r = 0.66, p = 0.03; BCa 95% CI: r = –0.24:0.9). The effect size of this relationship was found to be large for the complete sample (Cohen’s d = 8.35) and very large for the SA group alone (Cohen’s d = 45.65).

#### Amyloid deposition

No significant differences were observed in the AD-signature ROI composite for PIB SUVr between SA and C80 (1.25 ±0.24 vs. 1.32 ±0.25, p = 0.56) and in proportions of PIB positivity (30% SA vs. 30% C80 were PIB positive). Whole-brain PIB SUVr was observed to be correlated significantly with age in the entire sample (r = 0.51, p = 0.003; BCa 95% CI: r = 0.29:0.68), but did not reach statistical significance when calculated within groups (p > 0.05 for all groups). The relationship between whole-brain PIB SUVr and age also showed a large effect size (Cohen’s d = 8.29). The regional analysis revealed no statistically significant differences between groups in any of the previously described ROIs (p > 0.05 for all ROIs, Supplementary Table 1).

### 3.3. MRI results

#### Functional connectivity

The seed-based analysis revealed decreased connectivity between left sACC gyrus and left posterior cingulate cortex in the SA group as compared to that in the C80 group (cluster size = 1286 μL, p < 0.005, uncorrected; Fig. 3a). The cluster in the posterior cingulate cortex had a volume of 1286 μL and its peak coordinates were x=14.0, y=63.5, and z=4.0. After correction for multiple comparisons, a tendency of decreased functional connectivity was noted between the described regions in the SA group as compared to that in the C80 group.

**Figure 3.**
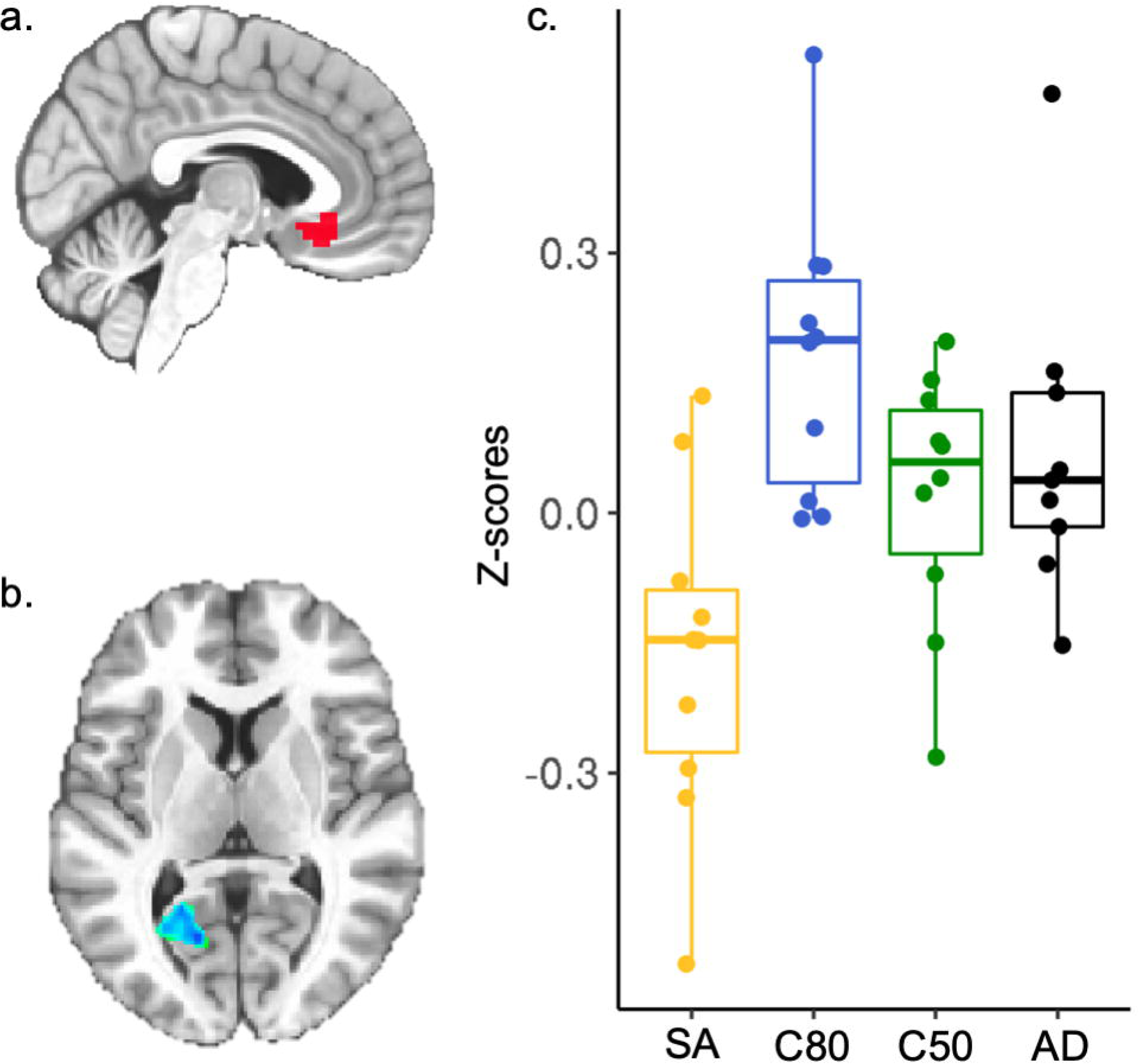
Seed-based functional connectivity. Decreased functional connectivity between left subgenual ACC region [in red–(a)] and left posterior cingulate cortex [in blue–(b)] in the SA group as compared to that in the C80 group. (c) Z-scores for the correlation between the left subgenual ACC seed and the cluster in the left posterior cingulate cortex in each group. SA – SuperAgers. C50 – Middle-aged controls. C80 – Age-matched controls. AD – Alzheimer’s disease.

**Figure 4.**
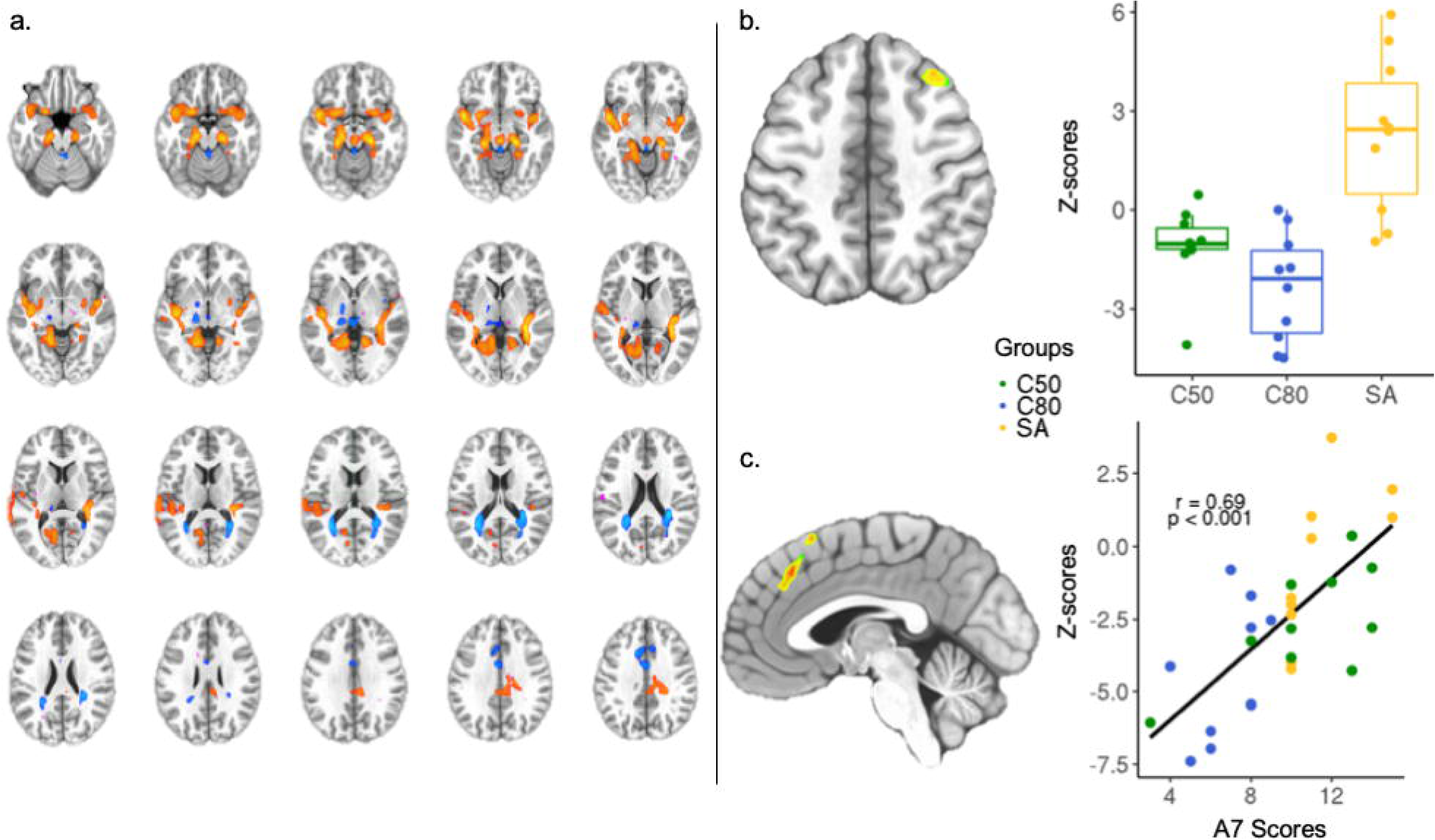
Independent component analysis of resting-state fMRI. (a) Identification of the Independent Component – 6 (Z-score = 4.033). (b) Independent component analysis showing right superior frontal gyrus, region within the IC–6 that distinguished SA from C80. (c) Regression analysis between delayed-recall memory scores and loading factors of IC–6 showing the right medial frontal gyrus and Pearson’s coefficient calculated between right medial frontal gyrus connectivity and the delayed-recall memory scores. SA – SuperAgers. C50 – Middle-aged controls. C80 – Age-matched controls. AD – Alzheimer’s disease.

ICA showed one component that significantly distinguished SA from the C80 group (IC–6), which involved abnormal functional connectivity of both medial, superior temporal, posterior cingulate and anterior cingulate areas (Fig. 4a). This network was similar to the Papez circuit. An increased functional connectivity of right superior frontal gyrus with the IC–6 (cluster size = 1414 μL, p < 0.005, uncorrected) was found for the SA group in comparison to the C80 group (Fig. 4b). A significant relationship (F = 9.464, p < 0.005 uncorrected) was also observed between left medial frontal gyrus in the IC–6 and the delayed-recall memory scores in the study sample, validated by a significant Pearson’s coefficient (r = 0.7, p < 0.001; BCa 95% CI: r = 0.46:0.83) (Fig. 4c). The AD group was not included in the ICA because it would bias the functional connectivity network analysis of other groups.

#### Cortical thickness and volume

Structural brain analysis did not reveal any difference between SA and C80 groups, when comparing cortical thickness and volume, after performing Monte-Carlo Z-null simulation correction. No statistically significant differences were found between total intracranial volume and bilateral mean cortical thickness between the SA and Control groups after correcting for multiple comparisons (p > 0.05, Monte Carlo Z-null simulation). However, a significant correlation between delayed-recall scores and the volume of the right hippocampus (r = 0.38, p = 0.03), though not with the left hippocampus (p = 0.18), was observed in the sample.

## 4. Discussion

A group of older adults with exceptional memory was identified and multimodal features of their brain areas supporting excellent memory function were examined. Selecting individuals with exceptionally high memory performance at an advanced stage of life may provide important biomarkers for memory maintenance through the aging process. The three main findings of this study were as follows: **Firstly**, SA showed subregions of anterior cingulate associated with brain activity similar to that of the middle-aged participants. **Secondly**, SA exhibited similar amyloid burden as their age-matched counterparts. **Thirdly**, in both hypothesis-driven and data-driven models of FC, SA presented altered functional connectivity in the frontal regions.

Measuring total years of education is very common, though it is a limited proxy of cognitive reserve in older adults. In contrast to the other studies on high-performing older adults, the cohort of SuperAgers in this study did not show a high level of education (Cook et al., 2017; Dekhtyar et al., 2017; Harrison et al., 2012). SuperAgers also showed a similar level of education as that of the normal agers, which eludes the assumption of increased cognitive reserve in this group.

Subregions of ACC were found to be associated with memory scores, independent of age and region volume, showing an even stronger association in SuperAgers. Specifically, the subgenual ACC area is a potential biomarker for memory maintenance in older adults. This region showed reciprocal connectivity with the hippocampus and many cortical and subcortical areas (Ongur, 2000). Though ACC is highly associated with regulation of emotions (Dunlop and Mayberg, 2014), there is growing evidence that this region is crucial for memory encoding (Schlichting and Preston, 2016) and remote memory retrieval (Ezzyat et al., 2018; Takashima et al., 2006). SuperAgers rely on higher brain activity in this region as compared to the normal agers, thereby supporting its significant relation with episodic memory performance. Furthermore, this region has a brain activity that is statistically similar to the middle-aged adults, indicating its uniqueness and youthfulness in high memory scores in older adults. It was hypothesized that both hippocampi and ventromedial prefrontal cortex are involved in memory consolidation differently. Over time, a decreased hippocampal activity and increased prelimbic prefrontal activity was noted during a remote memory retrieval task (Takashima et al., 2006). Also, sACC provides different representational aspects of memory to hippocampus, thus supporting prospective coding of information (Ezzyat et al., 2018; Guise and Shapiro, 2017). SuperAgers may present with a better memory differentiation than the normal agers, which ultimately improves memory consolidation.

Amyloid deposition in SuperAgers was similar to that of the normal agers, in this study. Recent studies on high-performing older adults presented inconsistent findings (Dekhtyar et al., 2017; Gefen et al., 2014; Lin et al., 2017), possibly due to high heterogeneity in the selection of the sample. Findings of this study are similar to those of previous studies on high-performing older adults (Dekhtyar et al., 2017), though the cohort included here are individuals of a higher age range only. Age threshold is important in the context of Alzheimer’s disease because of increased amyloid deposition rate seen in the non-demented older adults as they age (Jansen et al., 2015), substantiated by a positive correlation between age and amyloid burden in this study sample. Despite showing similar regional amyloid burden as their counterparts, SuperAgers exhibited preserved brain activity in frontal areas. This highlights a possible mechanism of better coping with brain pathology, specifically in SuperAgers’ brain as compared to that of cognitively normal older adults (Nyberg et al., 2012). A recent investigation indicated that an important dose-response effect reflects the association between amyloid burden and cognitive decline (Farrell et al., 2017), which suggests that the rate of accumulation is a better predictor of cognitive decline than a single threshold. Besides, it is known that cognitive decline is strongly associated with Tau deposition (Aschenbrenner et al., 2018; Braak et al., 2006; Schilling et al., 2016). Hence, further studies assessing longitudinal amyloid and tau deposition in SuperAgers may help elucidate this mechanism.

Reduced global functional connectivity is expected with aging, though some regions are more susceptible to age-related functional changes than the others (Dennis and Thompson, 2014). A decreased functional connectivity between left sACC and PCC was detected in SuperAgers as compared to that in the C80 group, which corroborates previous findings of decreased PCC and ACC functional connectivity in high performing adults (Lee et al., 2016). Compensatory local connectivity in older adults has been previously described in many models of cognitive aging (Cabeza et al., 2002; Davis et al., 2008; Park and Reuter-Lorenz, 2009). In general, these models try to elucidate age-related changes in brain connectivity, associated with an unexpected increase in regional connectivity. However, compensatory connectivity may actually be associated with less efficient networks in older adults (Morcom and Henson, 2018). Therefore, these compensatory shifts may not necessarily occur in a preserved brain architecture. SuperAgers also exhibited increased functional connectivity in frontal regions of the IC–6, a neural network involving hippocampal and anterior cingulate regions, similar to the Papez circuit (Papez, 1937). This network showed an increase in FC of right superior frontal gyrus in the SA group as compared to that of healthy older adults and middle-aged adults. Also, right medial frontal gyrus was significantly associated with episodic memory scores. Together, these findings suggest that frontal networks have increased connectivity in the Superaging process.

This study corroborates to the role of frontal areas in maintaining a more youthful memory ability in older adults. Preserved brain features observed in this study provide evidence for the theory of brain maintenance (Nyberg et al., 2012), as SuperAgers presented metabolic and functional brain features that were statistically similar to those of the younger group. The similarity of glucose metabolism between SuperAgers and middle-aged participants in left sACC indicates the avoidance of age-related hypometabolism typically seen in cognitively normal older adults (Knopman et al., 2014; Pardo et al., 2007). Frontal preservation was highlighted as fundamental in the preservation of memory functions during the aging process (Vidal-Piñeiro et al., 2018), which corroborates our findings. Besides, Dekhtyar et al. (2017) showed that executive functions, typically associated with the frontal activity, are maintained in high-performing older adults as compared to that in the normal performers.

Even though this study included small sample size, it is important to mention that the participants were selected through a very strict inclusion criterion of advanced age and exceptionally high cognitive scores. The sample size has been accounted as an important limitation as some outcomes did not survive the correction for multiple comparisons; thus, interpretation of the results should be carefully addressed. Even though the Apolipoprotein E (ApoE) status was unavailable for this sample, individuals without any type of family history of dementia or cognitive impairment were selected. Besides, a population-based study (Petersen et al., 2016) corroborated that PIB positivity is associated with cognitive decline, independent of APOE status. A longitudinal approach to this population is decisive to confirm whether all the described brain features are persistent or temporary. Further work is warranted to clarify the mechanisms of memory maintenance in a longitudinal and multimodal brain analysis.

Overall, these results authenticate the key role of anterior cingulate cortex in the exceptional memory ability of SuperAgers, even in the presence of amyloid deposition. Metabolic and functional changes of the subgenual and pre-subgenual areas may be essential in maintaining high memory performance in older adults. In particular, subgenual ACC is a potential biomarker of memory function in older adults.

## Acknowledgements

WVB, ELC and GR received a CAPES scholarship (PBE-DPM II, Programa de Excelência Acadêmica and PDES, respectively). JCC is funded by CNPq (Bolsa de produtividade de pesquisa). We would like to thank Luciana Borges Ferreira for support in the data acquisition of this study.

