## Supplemental material for "Neural correlates of exceptional memory ability in SuperAgers: A multimodal approach using FDG-PET, PIB-PET, and MRI" (from another version of this preprint: version 1, 2019-06-11)

**Supplementary material**

Table 1. Volumes of each selected ROI extracted by PMOD software.

|  | **SA** | **C80** | **C50** | **AD** | **SA vs. C80 p-values** |
| --- | --- | --- | --- | --- | --- |
| Right ACC | 4.61 (±0.45) | 4.51 (±0.50) | 5.24 (±0.55) | 3.72 (±0.80) | 0.98 |
| Left ACC | 4.46 (±0.71) | 4.84 (±0.97) | 5.31 (±0.54) | 3.75 (±0.68) | 0.67 |
| Right PCC | 4.38 (±0.84) | 4.49 (±0.79) | 4.62 (±0.45) | 4.02 (±0.98) | 0.99 |
| Left PCC | 4.71 (±0.95) | 4.95 (±0.89) | 5.25 (±0.51) | 4.13 (±0.98) | 0.92 |
| Right pACC | 0.43 (±0.06) | 0.39 (±0.05) | 0.50 (±0.08) | 0.35 (±0.09) | 0.51 |
| Left pACC | 0.67 (±0.07) | 0.63 (±0.17) | 0.75 (±0.06) | 0.50 (±0.10) | 0.78 |
| Right sACC | 0.75 (±0.08) | 0.65 (±0.07) | 0.79 (±0.10) | 0.56 (±0.10) | 0.06 |
| Left sACC | 0.81 (±0.10) | 0.84 (±0.17) | 0.85 (±0.12) | 0.71 (±0.14) | 0.93 |
| Right vACC | 0.59 (±0.06) | 0.52 (±0.05) | 0.65 (±0.08) | 0.45 (±0.09) | 0.13 |
| Left vACC | 0.74 (±0.08) | 0.73 (±0.16) | 0.80 (±0.09) | 0.61 (±0.11) | 0.99 |

SA – SuperAgers. C50 – Middle-aged controls. C80 – Age-matched controls. AD – Alzheimer’s disease. ACC – Anterior Cingulate Cortex. sACC – Subgenual ACC. pACC – Presubgenual ACC. vACC – Ventral ACC.

Table 2. PIB SUVr values for each selected Cingulate ROIs.

|  |  | **SA** |  | **C80** |  | **C50** | **AD** | **SA vs. C80 p-values** |
| --- | --- | --- | --- | --- | --- | --- | --- | --- |
| AD-signature VOI composite |  | 1.3 (±0.2) |  | 1.3 (±0.3) |  | 1.1 (±0.1) | 1.5 (±0.4) | 0.94 |
| Right ACC |  | 1.36 (±0.35) |  | 1.47 (±0.34) |  | 1.16 (±0.14) | 1.71 (±0.51) | 0.90 |
| Left ACC |  | 1.40 (±0.36) |  | 1.50 (±0.31) |  | 1.19 (±0.14) | 1.75 (±0.52) | 0.92 |
| Right PCC |  | 1.44 (±0.40) |  | 1.56 (±0.40) |  | 1.19 (±0.09) | 1.78 (±0.51) | 0.88 |
| Left PCC |  | 1.48 (±0.40) |  | 1.60 (±0.40) |  | 1.19 (±0.11) | 1.79 (±0.50) | 0.88 |
| Right pACC |  | 1.43 (±0.41) |  | 1.46 (±0.35) |  | 1.16 (±0.16) | 1.79 (±0.59) | 0.99 |
| Left pACC |  | 1.44 (±0.52) |  | 1.51 (±0.38) |  | 1.15 (±0.14) | 1.89 (±0.64) | 0.98 |
| Right sACC |  | 1.42 (±0.36) |  | 1.38 (±0.28) |  | 1.13 (±0.14) | 1.65 (±0.54) | 0.99 |
| Left sACC |  | 1.42 (±0.39) |  | 1.41 (±0.31) |  | 1.14 (±0.10) | 1.71 (±0.56) | 0.99 |
| Right rACC |  | 1.42 (±0.38) |  | 1.42 (±0.31) |  | 1.15 (±0.15) | 1.72 (±0.56) | 0.99 |
| Left rACC |  | 1.43 (±0.45) |  | 1.46 (±0.34) |  | 1.15 (±0.12) | 1.80 (±0.60) | 0.99 |

SA – SuperAgers. C50 – Middle-aged controls. C80 – Age-matched controls. AD – Alzheimer’s disease. ACC – Anterior Cingulate Cortex. sACC – Subgenual ACC. pACC – Presubgenual ACC. vACC – Ventral ACC.

Table 3. Correlation analysis between brain components, cognitive scores and age in the whole sample (AD group was not included in this analysis).

|  | RAVLT - A7 | RAVLT - Sum | Age |
| --- | --- | --- | --- |
| **FDG**  **(SUVr)** |  |  |  |
| Right pACC | 0.59 (0.3 - 0.76)*** | 0.62 (0.42 - 0.77)*** | -0.65 (-0.8 - -0.37)*** |
| Left pACC | 0.37 (-0.03 - 0.65) | 0.58 (0.26 - 0.79)*** | -0.67 (-0.8 - -0.43)*** |
| Right sACC | 0.49 (0.23 - 0.68)* | 0.65 (0.45 - 0.78)*** | -0.60 (-0.77 - -0.3)*** |
| Left sACC | 0.41 (0.12 - 0.62)* | 0.63 (0.4 - 0.76)*** | -0.60 (-0.78 - -0.35)*** |
| Right vACC | 0.55 (0.22 - 0.73)** | 0.65 (0.46 - 0.79)*** | -0.64 (-0.81 - -0.38)*** |
| Left vACC | 0.40 (0.05 - 0.65)* | 0.63 (0.36 - 0.79)*** | -0.66 (-0.8 - -0.43)*** |
| Right hippocampus | 0.46 (0.22 - 0.65)* | 0.54 (0.29 - 0.71)** | -0.57 (-0.75 - -0.31)*** |
| Left hippocampus | 0.47 (0.21 - 0.64)* | 0.57 (0.36 - 0.71)*** | -0.61 (-0.81 - -0.36)* |
| **PIB**  **(SUVr)** |  |  |  |
| AD-Signature ROI Composite | -0.13 (-0.57 - 0.33) | -0.11 (-0.50 - 0.22) | 0.51 (0.29 - 0.68)** |
| **FMRI**  **(Loading factors)** |  |  |  |
| IC-6 right medial frontal gyrus | 0.7 (0.46 - 0.83)*** | 0.49 (0.21 - 0.68)* | -0.07 (-0.41 - 0.17) |

Note: Shown are Pearson’s correlations and the 95% bias-corrected and accelerated confidence intervals for bootstrapped correlations. ACC – Anterior Cingulate Cortex. sACC – Subgenual ACC. pACC – Presubgenual ACC. vACC – Ventral ACC. RAVLT - Rey Auditory-Verbal Learning Test; A7 - delayed-recall memory score; SUM - Sum of the first 5 lists of the RAVLT. * p < 0.05; ** p < 0.005; *** p < 0.001

Table 4. Regression models adjusting for ROI volume and age.

|  | RAVLT - A7 | RAVLT - Sum |
| --- | --- | --- |
| **FDG (SUVr)** |  |  |
| Right pACC | **0.62 (10^-6^) [0.004]*** | **0.64 (10-6) [0.002]*** |
| Left pACC | 0.47 (0.0007) [0.38] | **0.54 (0.0001) [0.03]*** |
| Right sACC | 0.54 (0.0008) [0.07] | **0.63 (10-6) [0.003]*** |
| Left sACC | 0.40 (0.003) [0.19] | **0.52 (0.0001) [0.006]*** |
| Right vACC | **0.60 (10^-5^) [0.01]*** | **0.66 (10-6) [0.001]*** |
| Left vACC | 0.46 (0.0008) [0.25] | **0.56 (10-5) [0.009]*** |
| Right hippocampus | 0.39 (0.003) [0.09] | 0.41 (0.002) [0.05] |
| Left hippocampus | 0.46 (0.0008) [0.09] | **0.49 (0.0004) [0.04]*** |
| **PIB (SUVr)** |  |  |
| AD-Signature  ROI Composite | 0.26 (0.01) [0.7] | 0.27 (0.01) [0.4] |
| **FMRI**  **(Loading factors)** |  |  |
| IC-6 right medial frontal gyrus | **0.52 (10-5) [10^-6^]**** | **0.27 (0.01) [0.003]*** |

Note: Data is showed as follows: R-squared of the model (p-value of the overall model) [p-value of the cognitive score in the model]. *regression models in which the cognitive score showed p < 0.05 independently of overall R2. ACC – Anterior Cingulate Cortex. sACC – Subgenual ACC. pACC – Presubgenual ACC. vACC – Ventral ACC.

Figure 1. Subregions of Cingulate Cortex described in this study.

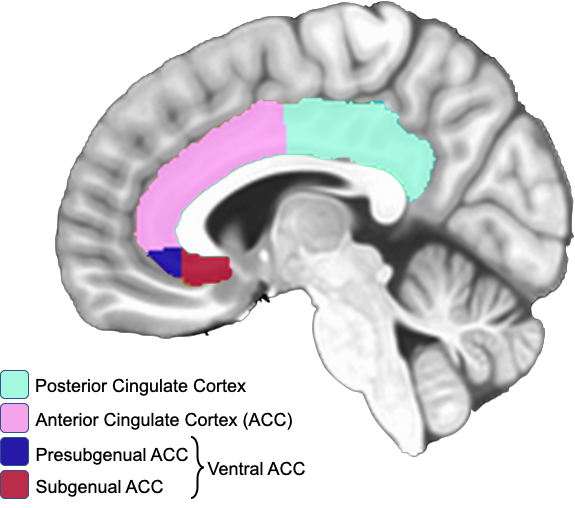

Figure 2. Alzheimer’s Disease-signature ROI SUVr comparison between groups. ACC – Anterior Cingulate Cortex. sACC – Subgenual ACC. pACC – Presubgenual ACC. vACC – Ventral ACC.

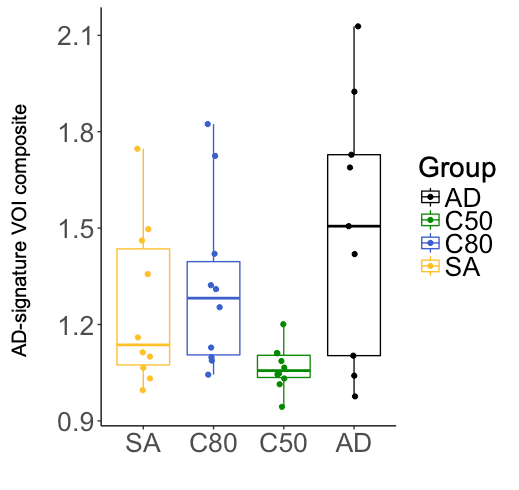
