## Supplementary material for "Neural correlates of exceptional memory ability in SuperAgers: A multimodal approach using FDG-PET, PIB-PET, and MRI": Highlights (from another version of this preprint: version 1, 2019-06-11)

- SuperAgers show increased metabolic activity in left subgenual anterior cingulate;
- Subareas of anterior cingulate are associated with delayed-recall memory scores;
- SuperAgers exhibit abnormal functional connectivity of frontal areas;
- Amyloid deposition is similar between SuperAgers and normal agers.
